# Enhancing additive series relay intercropping of grass pea (Lathyrus sativus L.) with low land rice (Oryza sativa L.) in North-Western Ethiopia: A farmer’s indigenous knowledge

**DOI:** 10.1101/2023.01.24.525334

**Authors:** Endeshew Assefa, Yayeh Bitew

## Abstract

In Ethiopia the facts of farmer’s indigenous knowledge based cropping system has been rarely investigated through research. A field experiment was conducted during 2021/2022 main cropping season at Fogera plain with the objective of examining the effect of additive series relay intercropping of grass pea with lowland rice on the grain yield of the component crops and production efficiency of the cropping system. The experiment consisted of a factorial combination of four seed proportion of grass pea (SPGP) (25%, 50%, 75% and 100% of the recommended seed rate of sole grass pea) relay intercropped with full seed rate of rice in four Rice: Grass pea spatial arrangements (SA) (1:1, 2:1, 3:1 and their mixed relay intercropping system). The treatments were arranged in Randomized Complete Block Design with three replications. Data on grain yield of the component crops were collected and analyzed using SAS-JMP-16 software. Results revealed that SPGP and SA had no significant effect on rice. The highest grain yield of grass pea was obtained when 25% SPGP was relay intercropped with rice in 1:3 SA (5.10 t ha^-1^). Maximum production efficiency in terms of total land output yield (9. 89 t ha^-1^) and land use efficiency (ATER =1.33), net benefit (33, 5176.79 Birr ha^-1^), marginal rate of return (21,428%) and positive monetary advantage index with lower competitive ratio was obtained when 50% SPGP was relay intercropped with rice in 1:3 SA. Thus, this mixture seems contributing in the development of sustainable crop production with a limited use of external inputs. Rice intercropping with other staple legume crops under residual soil moisture needs to be tested across locations and years to intensify the production efficiency and profitability of the cropping system.

## Introduction

Now-a-days self-sustaining, diversified, low-input, and energy-efficient agricultural systems like intercropping have been considered as the efficient way to achieve the sustainability in agriculture by many farmers, researchers, and policy makers’ worldwide [1–3]. A majority of the world’s poor farmers, particularly those located in tropical regions including Ethiopia, still depend for their food and income on multispecies agricultural systems, i.e. the cultivation of a variety of crops on a single piece of land [3, 4]. Among these agronomic concepts, intercropping systems (crop diversity in space and time) involving cereal-legume integrations is the common traditional cropping systems practiced worldwide and can be considered as a practical application of ecological principles including biodiversity conservation, plant interactions and other natural regulation mechanisms [5,6].This cropping system is assumed to have potential advantages in productivity, stability of outputs, resilience to disruption and ecological sustainability [6]. It also have resulted in increased farm production and profitability per unit land area [7, 8]. Researches showed that cereal-legume intercropping is widespread among smallholder farmers in sub-Saharan Africa due to over yielding resulted from (1) the ability of the legume to cope with soil erosion, (2) increase water use efficiency [9–11], (3) improve light interception [12], (4) improve nutrient use efficiency [10, 11] (5) controlling weeds, insect pest and diseases [13], (6) Lodging resistance to prone crops, (7) insurance against crop failure [14], (8) nitrogen transfer in cereal-legume intercropping systems [9], (9) residual effects of none legume-legume cropping system [15, 16]. Among the various cereal-legume intercropping systems/methods, relay intercropping of grass pea (as a supplementary crop) with low land rice (as a main crop) is one of common cropping system practiced by subsistence farmers in Northwestern Ethiopia [17].

Low land rice and grass pea are among the cereal and legume crops, respectively, grown in Fogera plain, Northwestern Ethiopia [17]. Currently, rice is considered as the millennium crop [18] while grass pea has been a neglected crop for many years in Ethiopia [19]. However, both crops are a food security crops mainly for subsistence farmers [17]. In descending order, *Amhara, Gambela, Southern Nations Nationalities and Peoples Region (SNNPR), Benshangul-Gumz, Oromia and Tigray* are the major rice producing regions in Ethiopia [20].The study area (Amhara region) accounts for 32% of the area coverage and 28.10% of the annual production in the country [21]. In Ethiopia grass pea is widely grown in the northwestern (58%), the central (16%) and the northeastern (13%) parts of the country. The northern and southeastern parts of the country account for the remaining 13% of grass pea area [22].

Rice as a main crop has been grown in both sole and intercropped with grass pea, while grass pea as a supplementary crop is grown in relay intercropped with rice in Fogera plain of northwestern Ethiopia [17]. Relay intercropping is a method of multiple cropping where one crop is seeded into standing second crop well before harvesting [14]. Some of the benefits of relay intercropping are: (1) optimizes system productivity and land use efficiency through growing two or more crops at same land in same cropping season (2) Improve soil quality and in turn increased the component crops yield due to the presence of N fixation in the system [3, 23].

Relay intercropping of grass pea with low land rice is an indigenous knowledge practiced by local farmers in rice producing areas of Northwestern Ethiopia. The most important reason that farmers used this kind of intercropping is to get additional yield from the supplementary crop (grass pea) and to improve the soil fertility of their cultivated land for the subsequent cropping season and in turn increased the productivity of low land rice [17]. Despite this benefit, this cropping system has been practiced only by the imagination of local farmers in which its suitability and profitability under research has not been yet investigated. Farmers practiced this cropping system in mixed additive series relay intercropping system (100% rice seed rate with ≤ 50% grass pea seed rate) without any row arrangement. The selection of proper sowing method, spatial arrangement and ratio with the application of various competition indices hence is crucial for adopting the right intercropping system in this environment. Therefore, the objective of this study were to determine the appropriate sowing methods, spatial arrangement and optimum seed proportion of grass pea in additive series relay intercropping of grass pea with low land rice for the highest land use efficiency and profitability with low competitive ratio.

## Material and methods

### Description of the study area

The field experiment was conducted during2021/2022 main cropping season at Fogera national rice research and training center, Northwestern Ethiopia. The experimental site is located at the Latitude of 11° 49’55” and Longitude of 37° 37’ 40 with an altitude from 1774 to 2516 m above sea level. The historical mean annual rainfall varies from 1216 mm to 1336 mm with the average minimum and maximum annual temperature of 12^0^c and 28^0^c, respectively [18,17]. Generally, the rainfall in the study area follows a dominantly unimodal distribution with the main peak in June to September, during which more than 80% of the annual rainfall is received. This experimental area is generally found in agro-ecological zone of moist *Wayena Dega* (mid land).The farming system of the study areas is characterized by 100% mixed crop-livestock systems [3].The total annual rain fall collected during the experimental year (2021/2022) was 1199 mm with the mean annual maximum and mean annual minimum temperature of 28.55°C and 18.65°C, respectively (Fig.1).

**Fig 1.**
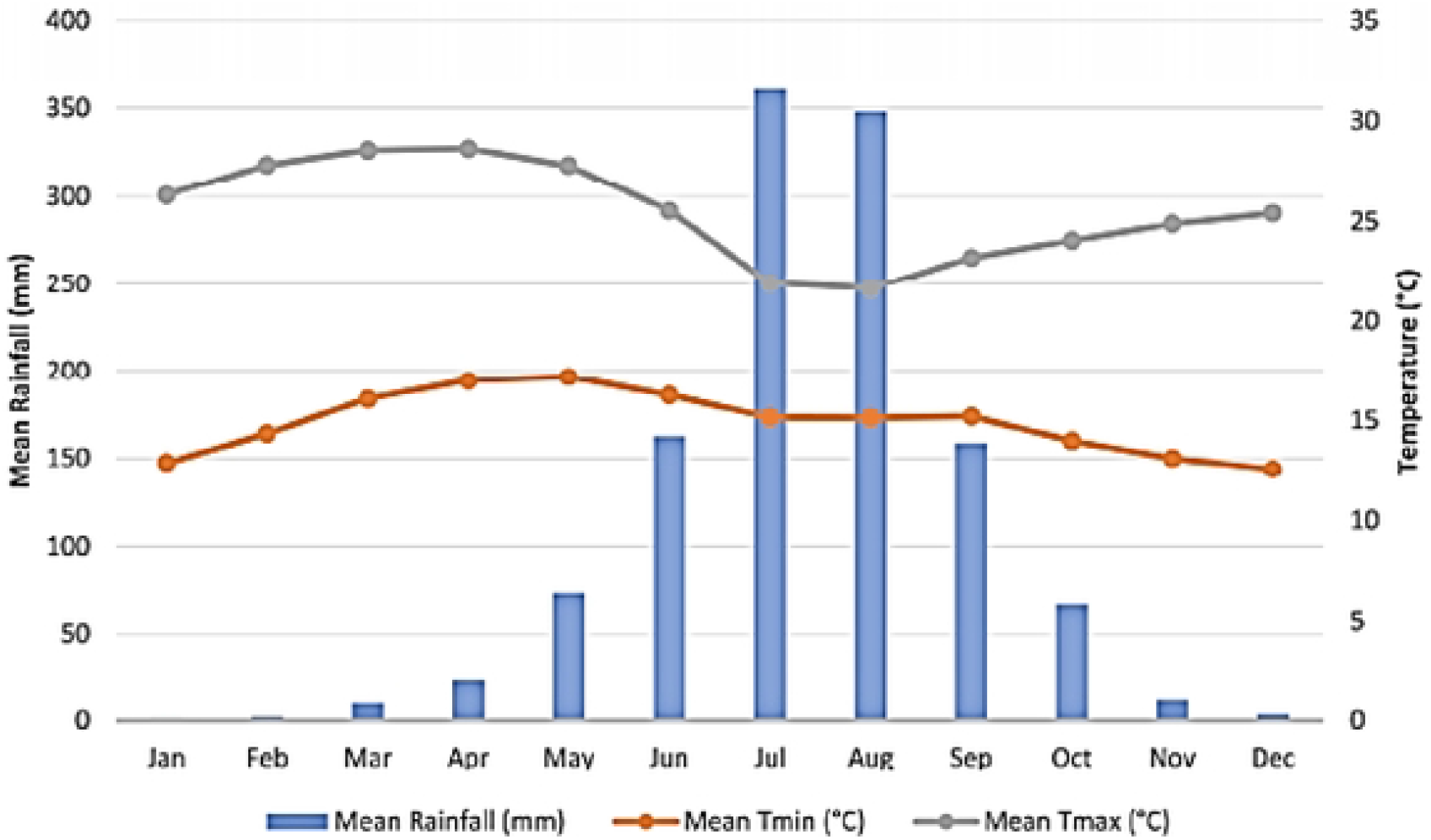
Monthly mean rainfall, maximum and minimum temperatures of the experimental site during 2021/2022 cropping season.

Immediately prior to implementing the experiment, composite soil samples for soil nutrient analysis were taken at five points diagonally at 0–20 cm soil depth for baseline information using soil auger. The collected soil samples were analyzed at Adet Agricultural Research center soil testing laboratory to determine some of the important physio-chemical properties of a soil. The soil texture, organic matter (OM), Organic carbon (OC), total nitrogen (TN), available phosphors (Ava.P), Soil reaction (pH), electrical conductivity (EC) and cation exchange capacity (CEC) were determined by Bouyoucos hydrometer method [24], Walkley and Black [25], Walkley and Black [26], Kjeldahl method [27], Bray-II method [28], pH meter [29], Rhoades method [30], Hesse method [31], respectively. Thus, based on the results of this soil laboratory analysis, the textural class of the soil found to be clay with a particle size distribution of 19% of sand, 15% of silt and 69% of clay. The soil pH value of the experimental site was 5.62 indicating that the soil reactions were found to be moderately acidic [32]. The other soil physio-chemical properties of the study area before planting is described in Table 1 below.

**Table 1.** Major soil physical and chemical characteristics of the experimental field before planting in 2021/2022 cropping season at Fogera national rice research and training centre, north-western Ethiopia.

| Soil properties | Units | Values | Rating | References |
| --- | --- | --- | --- | --- |
| **Particle size distribution** |  |  |  |  |
| **Sandy** | % | 19 |  |  |
| **Silt** | % | 15 |  |  |
| **Clay** | % | 69 |  |  |
| **Textural class** |  | Heavy Clay |  | [33] |
| **Soil chemical properties** |  |  |  |  |
| **PH** | - | 5.62 | Slightly acidic | [32] |
| **Organic Carbon** | % | 2.2 | Medium | [34] |
| **Organic Matter** | % | 2.61 | Medium | [34] |
| **Total Nitrogen** | % | 0.21 | Medium | [34] |
| **Available Phosphorus** | Ppm | 9.85 | Medium | [35] |
| **Electrical Conductivity** | ds/m | 0.05 | Low | [36] |
| **Cation Exchange Capacity** | C/molg | 57 | High | [32] |

### Experimental test material

The Improved cultivar low land rice *“Shaga”* and local grass pea variety were used in this experiment as a test crop. The selection criteria of *“Shaga”* variety is higher yield potential both in research and farmers field, resistance to cold, lodging and major rice diseases [37]. Local grass pea is wider adaptability and is default selected crop as it is dominantly grown in association with rice.

### Treatments and experimental design

The treatments consisted of four seed proportion of grass pea (SPGP)(25%, 50%, 75% and 100% sole grass pea seed rate) relay intercropped with full seed rate of sole rice in (i) three Rice (R): Grass pea (GP) spatial arrangements (SA) (row ratios) (1:1, 2:1 and 3:1) and (ii) rice (R): grass pea (GP) mixed intercropping (broadcast planting method). Based on the farmers practice, additive series intercropping experiment design was selected in which rice is the main crop component and grass pea is the supplementary (minor) crop component. Sole rice and sole grass pea were included as a comparison purpose (check) in an experiment. The experiment was laid out in a randomized complete block design (RCBD) with a total of 18 treatments (Table 2) in a factorial arrangement with three replications. The recommended seed rate of sole rice was 100kg ha^-1^ [18]. While the recommended seed rate of sole grass pea was 67kg ha^-1^ [38]. The gross and the net plot size were 3.4m x1.5m (5.1m^2^) and 1.8×1.5 (2.7m^2^) with a distance of 0.5m and 1m between adjacent plots and replications, respectively.

**Table 2.** Treatment combination used in additive series relay intercropping of grass pea with rice

| Trt No. | SPGP (%) | SA (R): (GP) | TCS |
| --- | --- | --- | --- |
| 1 | 25 | 1:1 | Row intercropping |
| 2 | 25 | 2:1 | Row intercropping |
| 3 | 25 | 3:1 | Row intercropping |
| 4 | 25 | MRI | Mixed intercropping |
| 5 | 50 | 1:1 | Row intercropping |
| 6 | 50 | 2:1 | Row intercropping |
| 7 | 50 | 3:1 | Row intercropping |
| 8 | 50 | MRI | Mixed intercropping |
| 9 | 75 | 1:1 | Row intercropping |
| 10 | 75 | 2:1 | Row intercropping |
| 11 | 75 | 3:1 | Row intercropping |
| **12** | 75 | MRI | Mixed intercropping |
| **13** | 100 | 1:1 | Row intercropping |
| **14** | 100 | 2:1 | Row intercropping |
| **15** | 100 | 3:1 | Row intercropping |
| **16** | 100 | MRI | Mixed intercropping |
| **17** | Sole rice | - | Row planting |
| **18** | Sole grass pea | - | Broadcasting |

Trt No., treatment number; SPGP (%), seed proportion of grass pea; SA, spatial arrangement; TCS, type of cropping system; GP, Grass pea; R, rice; MRI, mixed relay intercropping.

### Experimental procedure and cultural practices

The experiment was conducted under rain fed condition. Experimental plots were ploughed two times using tractor. The first plow was done during the dry season in January, the second plow was done at the end of May. Smoothing and leveling of the experimental plots was done mechanically at the same date that planting was done on 20 Jun 2021. Subsequently, rice planted in sole and intercropped with grass pea was sown with a seed rate of 100 kg ha^-1^ in 20cm drill row spacing. After 120 days of rice sowing date, one row of the associated supplementary crop (grass pea) was sown using drill row panting method after one, two and three rows of rice in 1:1, 2:1, 3:1 SP. All treatments containing rice-grass pea mixed intercropping was also done after 120 days of rice sowing date. Sole grass pea was planted in broadcasting planting method. For rice, a fertilizer rate 46kg N ha^-1^ in the form of UREA and 38 kg P_2_O_5_ in the form of NPS were applied [18]. None of the fertilizer types were applied on grass pea in all types of cropping systems. Full dose of P2O5 and 1/3 N were applied at rice planting, while remaining 2/3 of N were applied at flowering stage. All necessary cultural and agronomic practices were carried out uniformly for all plots as per the recommendation for each of the component crops.

### Data Collection and Measurements

#### Grain and straw Yield

To determine the grain and straw yield of both the component crops, above ground biomass in the net plot area were manually harvested, collected and sun dried with 27^0^C air temperature until the constant dry weight was attained. Dried above ground biomass was threshed and separated into grain (seed) and straw yields. Grain/seed yields were measured after adjusting the rice grain and the grass pea seed moisture content to 12.5% and 13%, respectively. The moisture correction factors was done by using the following formula [39, 40]:

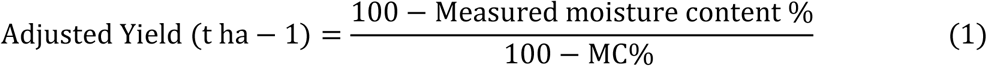

Where, MC% refers to moisture content of the component crops. Therefore, adjusted MC% of grain/seed yield = moisture correction factors X grain/ seed yield obtained from each plot.

#### Competition Ratio (CR)

Competition ratio defined as a measure of intercrop competition to measure the number of times by which one component crop is more competitive than the other [41]. Despite, many indices to compare the interspecific competition in an intercropping system, none of them effectively define the competition effect of the component crops [42] as with CR. Thus, competition between component crops in this cropping system was measured by the competition ratio [43] using the following formula.

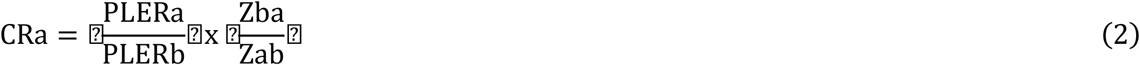

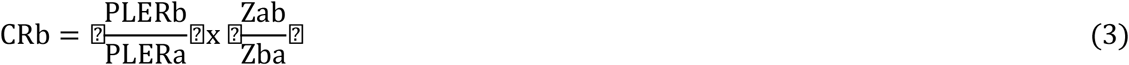

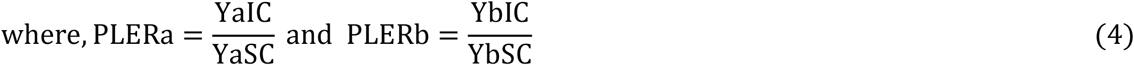

Where, CRa and CRb are a competitive ratio of crop *a* and *b*, respectively; PLERa and PLERb are a partial land equivalent ratio of crop *a* and *b*, respectively; and Zab and Zba are seed proportion of crop *a* in an intercropped with crop *b* and the seed proportion of crop *b* in an intercropped with crop *a*, respectively. YaIC and YbIC are yields of crop *a* and *b* in an intercropping, respectively, and YaSC and YbSC are the yield of crop *a* and *b* in sole cropping, respectively. If CRa is greater than one, it indicated that crop *a* was a competitor, while CRa less than one implied that crop *b* is suppressed crop *a* production.

### Determination of production efficiency

#### Land equivalent ratio (LER)

Land equivalent ratio (LER) is a measure for efficiency of land use in an intercropping experiment. It indicates the efficiency of intercropping for using the resources of the environment compared with mono-cropping [43, 44]. The LER was calculated using the formula outlined by Mead and Willy [44]:

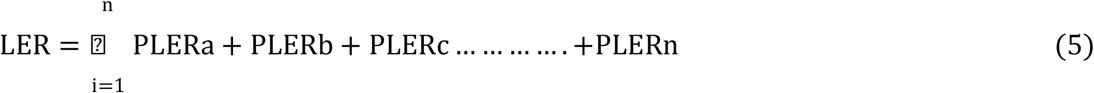

Where, n is the number of crops intercropped in the same areas in one growing season. The value of unity is the critical value. The null hypothesis was LER equal to one, indicating complementarity between two crops. When the LER was greater than one, intercropping favoured the growth and yield of the component crops. In contrast, when LER was lower than one, intercropping negatively affected the growth and yield of the component crops [19, 43].

#### Area Time Equivalent ratio (ATER)

Area time equivalent ratio provides more realistic comparison of the yield advantage of intercropping over monocropping in terms of time taken by component crops in the intercropping systems than LER. Area Time Equivalent ratio was calculated by formula developed by Hiebsch [45] as cited by Bitew et al. [3]:

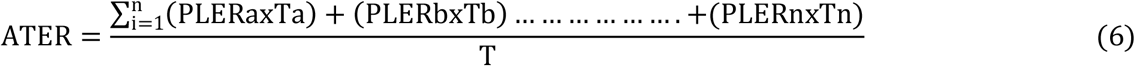

Where Ta, duration (in days) of crop *a* to reach maturity; Tb, duration (in days) of crop *b* to reach maturity and *T*, Total duration of the intercropping system (in days) to reach maturity. The interpretation of ATER is the same as with that of LER,

#### Rice Equivalent Yield (REY)

It is the conversion of crop yields into one form that allows us to compare the crops grown under mixed cropping, intercropping, or sequential cropping [46]. The total of intercrop crop *a* yield and converted crop *b* yield was used to compute crop *a* equivalent yield (*a*EY), which was then compared to the sole crop *a* yield. The conversion is done in the form of crop *a* equivalent yield by considering crop *b* yield and market price of crop *a* and crop *b* [47].

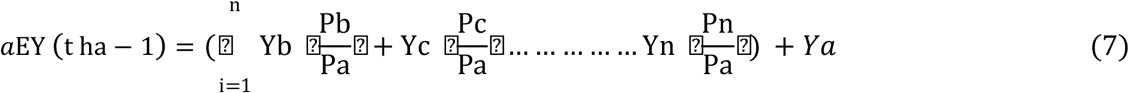

Where, a, b, c……n are type of crops grown in an intercropping; Y, yield; P, price

#### Total Land Output (TLOY)

Intercrop productivity was also assessed in terms of total land output Yield (TLOY). Total land output yield simply assess total production by a mixture, irrespective of densities or species combination [48]. Intercropped plots with greater TLOY values compared to monoculture showed yield advantage. TLO was computed as follows:

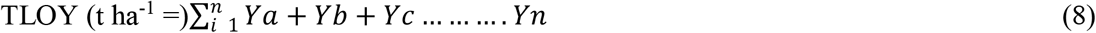

### Economic analysis

#### Monetary Advantage Index (MAI)

It was calculated to give some economic evaluation of intercropping as compared to sole cropping. The monetary advantage index (MAI) was calculated by the formula developed by Willey [49].

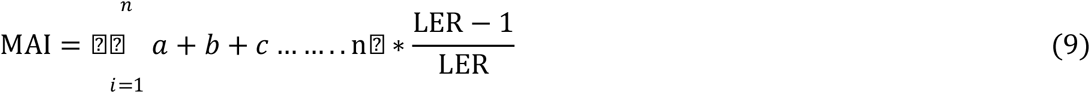

Where, a, b, c….n are component crops in an intercropping system.

### Partial budget analysis

To determine economically profitable treatment(s), partial budget analysis was performed following the [50] methodology. Seed (grass pea) and labor for grass pea planting and DAP fertilizer and fertilization for grass pea; harvesting, trashing and cleaning of both component crops costs were considered as the input costs. According to CIMMYT [50] only significant treatments are taken for Partial budget analysis to Thus, only grass pea straw and seed yield were considered as an income elements as rice straw and grain yields were non-significantly affected by the treatments. Other production costs such as labor for land preparation, weeding of rice, fertilizer application for rice were considered as fixed costs across treatments. All costs were calculated as average value of 2021 and 2022 on a per-hectare basis. The price of DAP was 17 birr kg^-1^, and the average labor cost was estimated 100 Ethiopian Birr (ETHB) man day^-1^. Based on the average local market prices in the months from June to February costs of grain and straw yield for grass pea were 44 and 2 ETHB kg^-1^, respectively The average costs of grain and straw yield in the same months for rice were 60 and 3 ETHB kg^-1^, respectively. Since yields of experimental plots are usually higher than that of farmers, yields of the treatments were adjusted down to 90% so as to reduce the yield gabs between experimental plots and farmers. Based on this information, total variable cost (TVC), gross benefit (GB); net benefit (NB) and marginal rate of return (MRR) were calculated as follows:

*Total variable cost (TVC):*

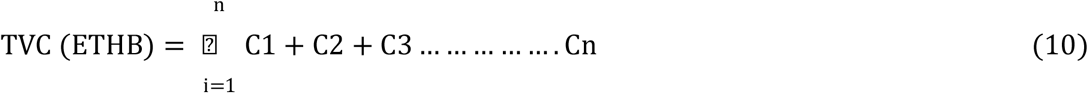
*Gross benefit:*

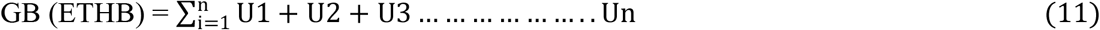
*Net benefit:*

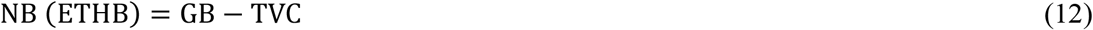

After treatments were arranged in ascending order by TVC value, treatments with high NB and lower TVC than the preceding treatment were selected for further analysis. Thus, treatments with a lower NB value and a greater TVC than the preceding were excluded. Selected treatments were subjected to marginal rate of return (MRR) analysis, which was calculated by the following formula:

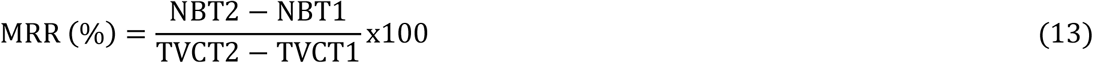

Where T2 and T1 are consecutive treatments (T) arranged in ascending order based on their TVC after excluding treatments with low NB and high TVC.

### Data Analysis

Quantitative data of the component crops from the experimental field was entered to Microsoft Office Excel. The same software was used for data management. Data analyses for the component yields were conducted using Statistical Software [51]. Before analysis, data were checked for normal distribution following the scatter plot technique. Data were analyzed with grass pea seed proportion and spatial arrangement as fixed effects, and replication as random effects. First, data analysis of the yield of the component crops in all treatments (18) were subjected to analysis of variance (ANOVA) following single degree of freedom orthogonal contrasts method. When there were significant difference between treatments at any probability level, mean separation was done using the Tukey-Kramer HSD test. However, if there were no significant difference (*P>0.05*) between all treatments (18), the data analysis of the same data in sixteen treatments (excluding the sole crops) were subjected to analysis of variance (ANOVA) following the same procedure. Mean separation was done using the same test, when there were significant difference between sixteen treatments at any probability level. Regression analysis was carried out to examine the relationship between production efficiency (total land output and rice equivalent yield) with a seed proportion and spatial arrangement.

## Result and discussion

### Grain (seed) yield of the component crops

Analysis of variance revealed that rice grain yield was not significantly (*P>0.05*) affected by the main effects of seed proportions of grass pea (SPGP) and spatial arrangements (SA) and by their interaction effects (Figure 2). Rice grain yield in this experiment ranges from 4.25 t ha^-1^ when 100% seed rate of grass pea was relay intercropped with rice in 1:3 SA to 5.32 t ha^-1^ when 50% seed rate of grass pea was relay intercropped with rice in 1:2 SA. This result is in conformity with the results of Yayeh and Fikremariam [17] who reported that all the agronomic attributes of rice were not significantly (*P>0.05*) affected by the main effects of seed proportions of chick pea and spatial arrangements and by their interaction effects. This might be due to early planting of rice in relay intercropping system takes an advantage in peak resource demands for nutrients, water, and sunlight for all treatments. The competition ratio (CR) of rice in grass pea-rice intercropping as indicated in Figure 3 confirms this remarks. Moreover, Bitew et al. [3] and Isaac et al. [52] also revealed that all the yield and yield components of maize were not significantly (*P>0.05*) affected by the main effects of seed proportion of haricot bean and spatial arrangement and by their interaction.

**Fig 2.**
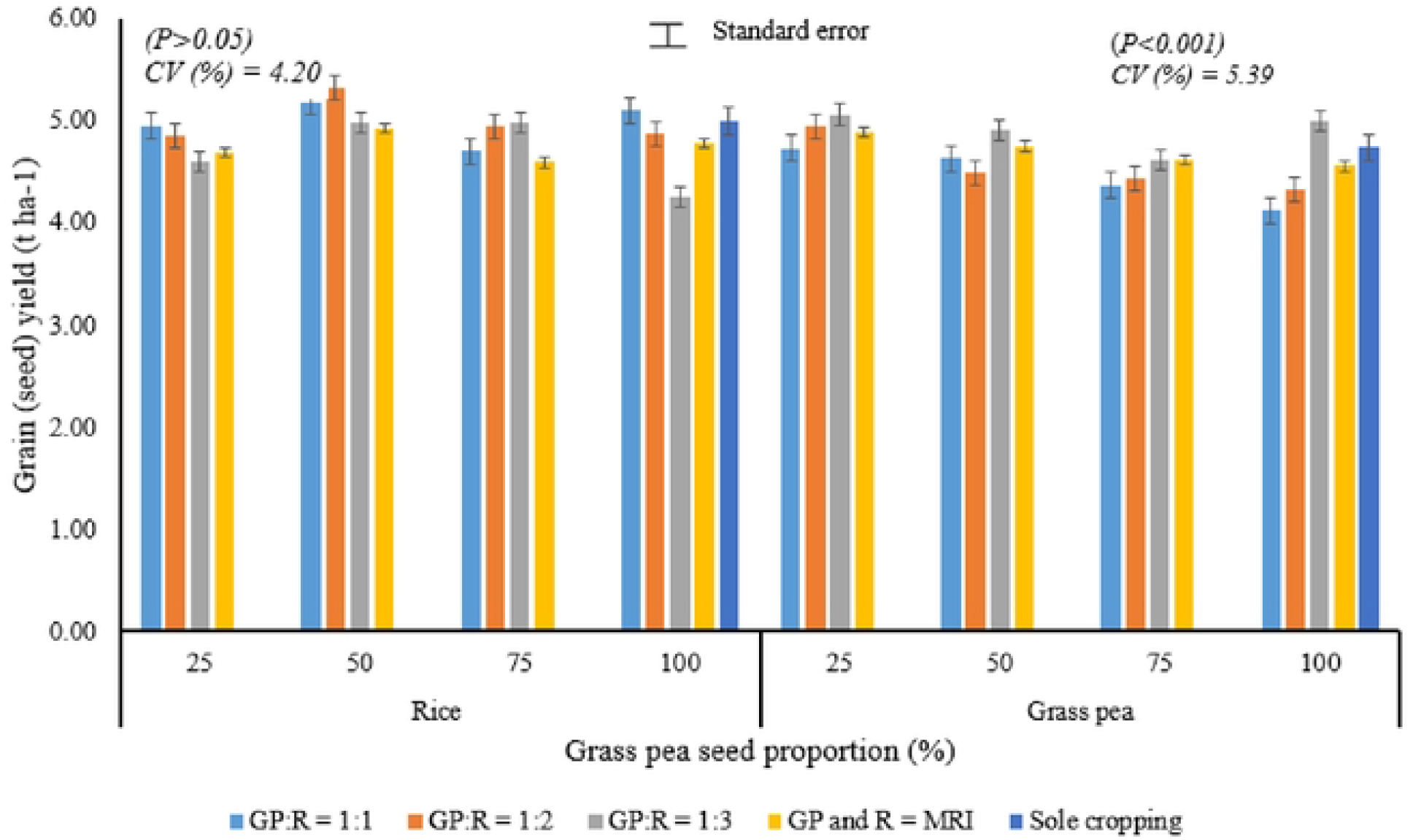
Iinteraction effects of SPGP (%) and (SA) in additive series relay intercropping of grass pea (GP) with rice (R) on the component crops economic yield at Fogera national rice research and training centre. GP:R = 1:1, 1:2, 1:3 indicating that grass pea was planted after one, two and three rows of rice, respectively; GP and R = MRI indicating that grass pea was mixed relay intercropped with rice.

**Fig 3:**
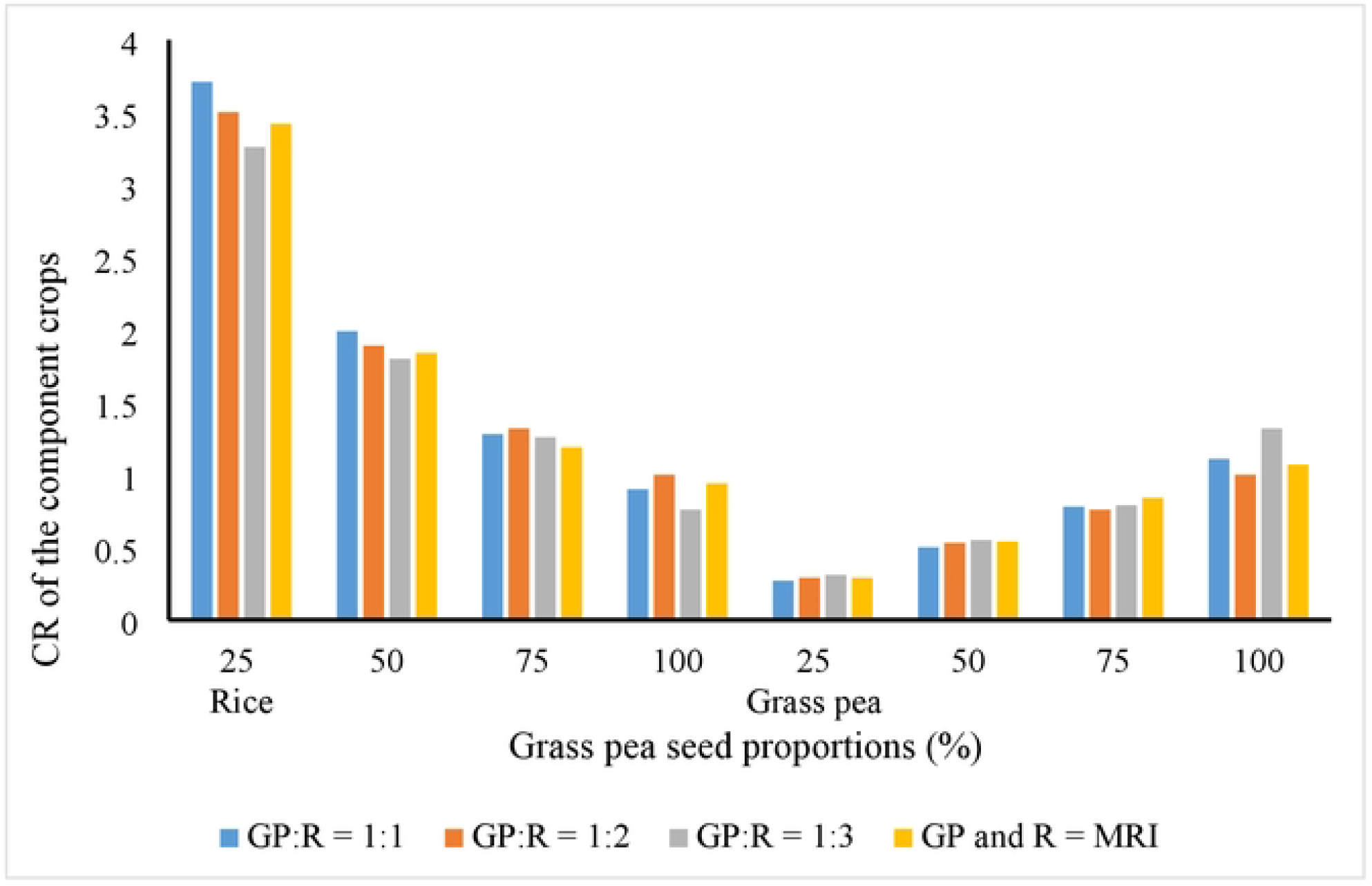
Competitive ratio of the component crops in grass pea-rice relay intercropping.

On the other hand, results revealed that grain yield of grass pea was significantly (*P<0.05*) affected by the main effect of seed proportion of grass pea and spatial arrangements and by their interaction (Figure 2). Maximum grain yield (5.06t ha^-1^) was obtained when 25% SPGP was relay intercropped with rice in 1:3 SA. This cropping system gave 7% higher grass pea seed yield over the sole cropping system. However, it is statistically similar effect with the treatments treated with the same SPGP with 1:2 SA (4.94 t ha^-1^) and mixed relay intercropping (MI) (4.89 t ha^-1^), 50% (4.90 t ha^-1^) and 100% (4.99 t ha^-1^) SPGP was relay intercropped with rice in 1:3 SA. While, minimum grain yield was obtained when 100% SPGP was relay intercropped with rice in 1:1 SA (4.12 t ha^-1^) (Figure 2). The higher grain yield in the former cropping system might be due to lower plant population resulted from the wider plant spacing between rows of grass pea that caused lower inter- and intra- plant competition. This result contradicted with the results of Banik et al. [53], who observed that the chick pea yield in wheat-check pea mixture significantly lower than sole cropped check pea.

#### Competition ratio (CR)

Rresults revealed that the CR of rice was greater than one except when 100% seed rate of grass pea was relay intercropped with rice in all SA while the reverse is true for grass pea (Figure 3). This might be due to grass pea was very low growth resource use (mainly) during the cogrowth with rice. In other words CR of grass pea was lower than CR of rice indicating that rice is the competitive crop in terms of growth resource use, especially light as compared to grass pea (rice created shading effect and in turn affected the growth and yield of grass pea) (Figure 3). This result was in agreement with the findings of Yayeh et al. [54], who stated that competitive ratio of wheat was higher than local lupine in local lupine- cereal intercropping system.

#### Land use efficiency

Results revealed that partial land equivalent ratio (PLER) of rice and grass pea in all cropping systems were greater than 0.5 (Figure 4) indicating that there was null or very low growth resource competition between the component crops during their co-growing period which supports the CR values. This result is inconsistent with the findings of Chen et al. [55], who reported that in sorghum–cowpea intercropping system PLER for cowpea and sorghum was lower and higher than 0.5, respectively, which indicated that a disadvantage for cowpea and an advantage for sorghum in an intercropping. Although, the combined yield advantage in terms of land equivalent ratio (LER) indices (Figure 4) was greatest when 50% SPGP was relay intercropped with rice in 1:3 SA (1.99), LER values of all intercropping system was sufficiently greater than one (Figure 4) indicating that growing the component crops in an intercropping was advantageous than growing the component crops individually. This result indicates that 0.83-0.99 more hectare of land is required by the sole cropping system to equalize the yield obtained from the intercropping system or about 83%-99% yield advantage was obtained from the intercropping over the sole cropping. Parallel to this result, Egbe et al. [56] demonstrated that most intercropping systems were more effective in utilization of environmental resources for growth and yield formation over sole cropping. Similar results were reported in intercropping of pea and barley [57], wheat and field bean [58] and maize and faba bean [59]. Regardless of the SA, as the SPGP increased from 25 to 100% the TLOY of the cropping systems decreased by 4.63%. This yield reduction could be mainly due to intense competition in soil residual moisture as the grass pea plant population increased.

**Fig 4.**
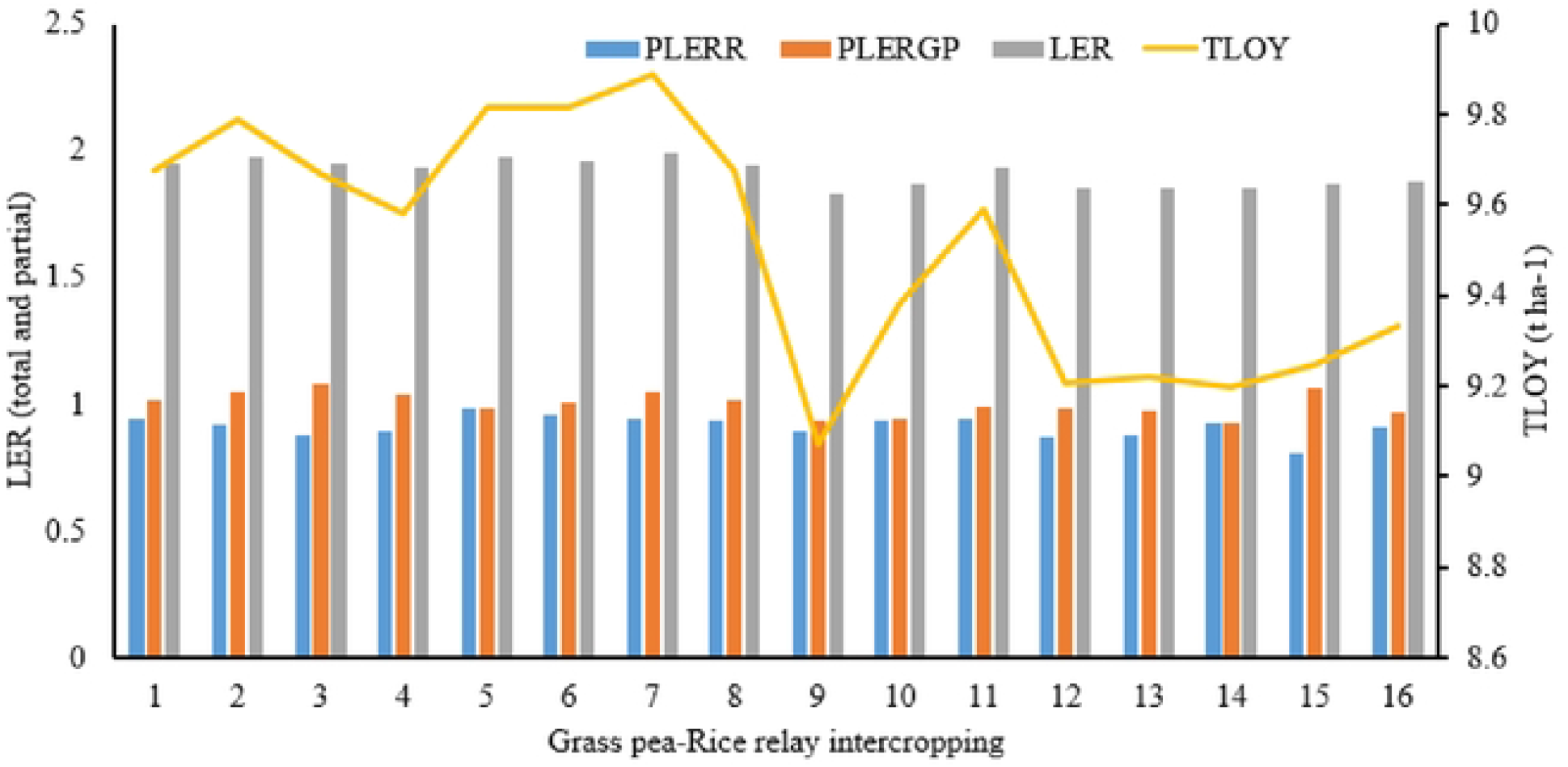
Land equivalent ratio (LER) (total and partial) and total land output yield (TLOY) in additive series relay intercropping of grass pea with rice. Note: 1 = 25%+1:1, 2 = 25%+2:1, 3 = 25%+3:1, 4 = 25%+M, 5 = 50%+1:1, 6 = 50%+2:1, 7 = 50%+3:1, 8 = 50%+M, 9 = 75%+1:1, 10 = 75%+2:1, 11 = 75%+3:1, 12 = 75%+M, 13 =100%+1:1, 14 = 100%+2:1, 15 =100%+3:1, 16 = 100%+M, PLERR = partial land equivalent ratio of rice, PLERGP = partial land equivalent ratio of rice grass pea.

The LER doesn’t consider the duration of the crops in the field and it is based on the harvested products, and not on desired yield proportion of the component crops [60]. Moreover, the choice of sole cropped yield for standardizing mixture yield in the estimation of LER is not clear [49]. Therefore, ATER provides more realistic comparison of the yield advantage of intercropping over sole cropping in terms of variation in time taken by the component crops of different intercropping systems [54]. In all treatments, ATER values are lesser than LER values (Table 3 and Figure 4) which indicated that over estimation of resource utilization in LER than ATER. Regardless of the values, parallel to LER, higher ATER was obtained when 50% SPGP was relay intercropped with rice in 1:3 (1.33) and 1:2 (1.32) SA with the yield advantage of 33% and 32%, respectively as compared to sole cropping (Table 3). This could be due to the reason that intercropping systems can actually give more efficient total resource exploitation and greater overall production than sole crops [61]. Similarly, compared with corresponding sole crops, yield advantages have been recorded in many cereal-legume intercropping systems, including groundnut-cereal fodders [62], barley-pea [55], and faba bean-barley [63], bean-wheat [64], lupine- finger millet [54], and finger millet-haricot bean [3].

**Table 3.** Effects of seed proportion of grass pea and spatial arrangement on the production efficiency (ATER, and LER) in rice-grass pea relay intercropping system at Fogera national rice research and training centre in 2021/2022

| Treatments combination |  | Partial LER |  | ATER | REY (t ha<sup>-1</sup>) | MAI |
| --- | --- | --- | --- | --- | --- | --- |
| SR | - | 1.000 |  | 1.000 | 5.00 | 1 |
| SGP | - |  | 1.000 | 1.000 | 3.48 | 1 |
| 25% | 1:1 | 0.94 | 1.01 | 1.28 | 4.96 | 1466.27 |
|  | 2:1 | 0.92 | 1.05 | 1.31 | 5.17 | 1516.93 |
|  | 3:1 | 0.88 | 1.08 | 1.28 | 5.29 | 1498.93 |
|  | MRI | 0.89 | 1.04 | 1.28 | 5.12 | 1458.2 |
| 50% | 1:1 | 0.98 | 0.99 | 1.31 | 4.86 | 1375.13 |
|  | 2:1 | 0.96 | 1.01 | 1.32 | 4.72 | 1465.6 |
|  | 3:1 | 0.94 | 1.05 | 1.33 | 5.14 | 1535.43 |
|  | MRI | 0.93 | 1.01 | 1.30 | 4.98 | 1365.4 |
| 75% | 1:1 | 0.90 | 0.94 | 1.22 | 4.60 | 1270.67 |
|  | 2:1 | 0.93 | 0.94 | 1.26 | 4.66 | 1342.63 |
|  | 3:1 | 0.94 | 0.99 | 1.29 | 4.85 | 1707.53 |
|  | MRI | 0.87 | 0.98 | 1.24 | 4.85 | 1333.27 |
| 100% | 1:1 | 0.88 | 0.98 | 1.25 | 4.32 | 1274.4 |
|  | **2:1** | 0.93 | 0.93 | 1.24 | 4.56 | 1295.9 |
|  | **3:1** | 0.81 | 1.07 | 1.22 | 5.23 | 1395.4 |
|  | **MI** | 0.91 | 0.97 | 1.25 | 4.79 | 1353.03 |

SPGP, Seed proportion of grass pea; SA, Spatial Arrangement; R, Rice; GP, Grass pea; ATER, Area time equivalent ratio; REY, Rice equivalent yield; MAI, monitory advantage index; MRI, mixed relay intercropping; CRGP, competition ratio of grass pea

The rice equivalent yield for different rice-grass pea relay intercropping system was higher when 25% of SPGP was relay intercropped with rice in 1:3 (5.29 t ha^-1^) followed in 1:2 (5.17 t ha^-1^) SA (Table 3). This was due to higher yield from the intercropped grass pea component and higher price of grass pea in the market. However, in grass pea – rice relay intercropping system, planting 50% SPGP in 1:3 SA ranked first with the highest TLOY (9.89 t ha^-1^) followed by when 50% SPGP in 1:2 SA (9. 81tha^-1^) (Figure 4). This finding further confirm the reports of Banik et al. [53] who reported that intercropping of wheat with chick pea gave higher total land output yield than sole cropped wheat and chickpea.

#### Economic analysis

The monetary advantage index (MAI) values were positive in all intercropping systems in the present study (Table 3) indicating that this intercropping systems were more profitable as compare to sole cropping. These results suggest that intercropping could improve the system’s productivity and increase the income for smallholder farmers. Monetary advantage index was higher in rice-grass pea relay intercropping when 75% of SPGP was relay intercropped with 1:3 SA (1707.53). These results imply that it was more economically viable and implying that the general suitability of grass pea as relay intercropping with rice when 75% SPGP planted in 1:3 SA in the study area. This result in line with the results of Yayeh et al. [54] and Tenaw et al. [65] who reported that intercropping of lupine with wheat and finger millet and barley-faba bean, respectively gave more economic returns and higher MAI as compared to sole cropping. In contrary, the highest net benefit was recorded when 50% (335,176.79 ETB ha^-1^) followed by 25% (330,718.47 ETB ha^-1^) SPGP was relay intercropped with rice in 1:3 SA with a marginal rate return (MRR) 21438% and 9551%, respectively. The lowest net benefit (294,659.52 ETB ha^-1^) was recorded when 75% SPGP was relay intercropped with rice in 1:1 SA (Table 4).

**Table 4.** Net benefit (NB) and marginal rate of return (MMR) in rice-grass pea relay intercropping as affected by seed proportion of grass pea and spatial arrangement at Fogera national rice research and training center in 2021/2022.

| **Treatments** | **TVC (ETB)** | **Gross benefit (ETB)** | **Net benefit (ETB)** | **MRR (%)** | **Dominance** |
| --- | --- | --- | --- | --- | --- |
| **25%+1:1** | 1191.12 | 320407 | 319215.9 | - |  |
| **25% +2:1** | 1278.88 | 328877 | 327598.1 | 9551 |  |
| **25%+3:1** | 1377.53 | 332096 | 330718.5 | 3163 |  |
| **25%+MI** | 1497.12 | 321830 | 320332.9 |  | D |
| **50%+1:1** | 2257.8 | 321207 | 318949.2 |  | D |
| **50%+2:1** | 2345.56 | 317660 | 315314.4 |  | D |
| **50%+3:1** | 2438.21 | 337615 | 335176.8 | 21438 |  |
| **50%+MI** | 2557.8 | 321920 | 319362.2 |  | D |
| **75%+1:1** | 3324.48 | 297984 | 294659.5 |  | D |
| **Sole rice and grass pea** | 3390.88 | 328312 | 324921.1 |  | D |
| **75%+2:1** | 3412.24 | 308611 | 305198.8 |  | D |
| **75%+3:1** | 3504.89 | 317462 | 313957.1 |  | D |
| **75%+MI** | 3624.48 | 307611 | 303986.5 |  | D |
| **100%+1:1** | 4390.88 | 300751 | 296360.1 |  | D |
| **100%+2:1** | 4478.64 | 302870 | 298391.4 |  | D |
| **100%+3:1** | 4571.29 | 320703 | 316131.7 |  | D |
| **100%+MI** | 4690.88 | 311847 | 307156.1 |  | D |

Where, TVC=Total variable cost, MRR=Marginal rate of return, ETBB=Ethiopian birr

## Conclusion

The study confirmed that seed proportion of grass pea and spatial arrangement had no significant effect on rice crop in additive series relay intercropping of grass pea with low land rice. The highest grain yield of grass pea was obtained when 25% SPGP was relay intercropped with rice in 1:3 SA. However, maximum production efficiency in terms of TLOY and land use efficiency, NB, MRR and positive MAI with lower CR was obtained when 50% SPGP was intercropped with the full seed rate of rice in 1:3 SA indicating that this cropping system is far better production system as compared to what farmers currently used (mixed intercropping system). Thus, this mixture seems contributing in the development of sustainable crop production system with a limited use of external inputs. The following research gaps were suggested for further research (i) the experiment need to be repeated across locations and years as the experiment was conducted in specific area and in one growing season; (ii) the effect of grass pea on soil fertility needs to be investigated as rice (the main crop) -grass pea (a supplementary crop) relay intercropping is the dominant cropping system in the study area; (iii) rice intercropping with other staple legume crop needs to tested to intensify the production efficiency and profitability of the cropping system.

## Acknowledgements

This research was done in collaboration with Ethiopian ministry of education. We wish to thank all people involved during data collection, writing and publication of this paper and particularly the individual farmers who rent us their farm land.

## Author Contributions

Conceptualization, Data curation, Formal analysis, Investigation, Methodology, Resources, Software, Supervision, Validation, Visualization, Writing – original draft, Writing – review & editing.

## Data Availability Statement

All relevant data are within the paper.

## References

1. Altieri M.A. Biodiversity and Pest Management in Agroecosystems. Food Products Press, New York, USA; 1994.

2. Jackson L. E., Pascual U. and Hodgkin T. Utilizing and conserving agro biodiversity in agricultural landscapes. Agriculture, ecosystems and environment. 2007; 121(3): 196–210.

3. Bitew Y., Alemayehu G., Adgo E., Assefa A. Competition, production efficiency and yield stability of finger millet and legume additive design intercropping. Renewable Agriculture and Food Systems 2912; 36: 108–119.

4. Malezieux E., Crozat Y. and Dupraz C. Mixing plant species in cropping systems: concepts, tools and models. Agronomy for Sustainable Development 2009; 29: 43–62.

5. Andersen M. K. Competition and complementarity in annual intercrops-the role of plant available nutrients (Doctoral dissertation, Samfundslitteraur Grafik, Frederiksberg, Copenhagen); 2005.

6. Thierfelder C., Cheesman S., Rusinamhodzi L. Benefits and challenges of crop rotations in maize-based conservation agriculture (CA) cropping systems of southern Africa. Int. J. Agric. Sustain. 2012; 1–17.

7. LaMondia J. A., Elmer W.H., Mervosh T.L. and Cowles R.S. Integrated management of strawberry pests by rotation andintercropping. Crop Protection 2002; 21: 837–846.

8. Yayeh B., Getachew A., Enyew A. & Alemayehu A. Boosting land use efficiency, profitability and productivity of finger millet by intercropping with grain legumes. Cogent Food & Agriculture 2019; 5:1.

9. Lithourgidis A.S., Dordas C.A., Damalas C.A., Vlachostergios D.N. 2011. Annual intercrops: an alternative pathway for sustainable agriculture. AJCS. 2011; 5(4): 396–410.

10. Li L. Z. and Fusuo Z. Crop Mixtures and the Mechanisms of Over yielding. In: Levin S.A. (eds.). Encyclopedia of Biodiversity 2013; 2: 382–395.

11. Matusso J. M., MugweJ.N. & Mucheru-Muna M. Potential role of cereal-legume intercropping systems in integrated soil fertility management in smallholder farming systems of Sub-Saharan Africa. Research Journal of Agriculture and Environmental Management 2014; 3(3): 162–174.

12. Seran T. H. and Brintha I. Review on maize based intercropping. Journal of agronomy 2010; 9(3): 135–145.

13. Willey R. W. Resource use in intercropping systems. Agricultural water management 1990; 17(1-3): 215–231.

14. Yayeh B. and Merkuz A. Conservation Agriculture Based Annual Intercropping System for Sustainable Crop Production: A review. Indian Journal of Ecology 2018; 46 (4): 235–249.

15. Ofori, F., Stern W.R. Cereal-legume intercropping systems. Adv. Agron. 1987; 40,: 41–90.

16. Yusuf A.A. Iwuafor N.O. Abaidoo R.C. Olufajo O.O. and Sanginga N. Effect of crop rotation and nitrogen fertilization on yield and nitrogen efficiency in maize in the northern Guinea savanna of Nigeria. African Journal of Agricultural Research 2009; 4 (10): 913–921.

17. Yayeh B. Fekremariam A. Rice (Oryza Sativa) and Chickpea (Cicer aritinum L) Relay Intercropping Systems in an Additive Series Experiment in Rain Fed Lowland Ecosystem of Fogera Vertisols. Journal of Biology, Agriculture and Healthcare2014; 4 (27): 199–204

18. Tilahun T. and Zelalem T. Review of rice response to fertilizer rates and time of nitrogen application in Ethiopia. International Journal of Applied Agricultural Sciences 2019; 5(6): 129–137.

19. Yayeh B. Integrating legumes for sustainable intensification of finger millet (Eleusine coracana) production in north western Ethiopia, PhD thesis, Bahir Dar University, Bahir Dar, Ethiopia; 2020.

20. Dawit A. Rice in Ethiopia: Progress in Production Increase and Success Factors 6th CARD General Meeting Ethiopia institute of agriculture research; 2015.

21. Kirub A., Alemu D., Shiratori K. and Assefa K. Challenges and opportunities of rice in Ethiopian agricultural development; 2011

22. Wolde-Amlak A., Asgelil D. B.., Regassa E., Wasie H.and Yeshanew A. Grass Pea (Lathyrus sativus) Research and Ms Production Potential in Ethiopia. A paper presented at the 3rd Triennial Colloquium of IN ILSEL, Dhaka. Bangladesh, 30 November-3 December 1991; 1991

23. Jabbar A., Ahmad R., Bhatti I. H., Aziz T. and Nadeem M. Residual Soil Fertility as Influenced by Diverse Rice-based Inter/Relay Cropping Systems. International Journal of Agriculture and Biology 2011; 13(4): 477–483

24. Bouyoucos G. J. Hydrometer method improved for making particle size analyses of soils 1. Agronomy journal 1962; 54(5): 464–465.

25. Walkley A. and Black I. A. An examination of the Degtjareff method for determining soil organic matter, and a proposed modification of the chromic acid titration method. Soil science 1934; 37(1): 29–38.

26. Walkley A. and Black C.A. An examination of the Degtjareff methods for determining soil organic matter and proposed modification of the chromic acid titration methods. Madison (WI): American Society of Agronomy; 1954.

27. Bremner J. M. and Mulvaney C. S. Total nitrogen. In. AL Page (ed.) Methods of Soil Analysis. Part 2. Chemical and Microbiological Methods; 1982.

28. Bray R. H. and Kurtz L. T. Determination of total, organic, and available forms of phosphorus in soils. Soil science 1945; 59(1): 39–46.

29. Chopra S. L. and Kanwar J. S. Analytical Agricultural Chemistry 1976; 245–298. Ludhiana.

30. Rhoades J. D. Salinity: Electrical conductivity and total dissolved solids. Methods of soil analysis: Part 3 Chemical methods 1996; 5: 417–435.

31. Hesse P. R. and Hesse P. R. A textbook of soil chemical analysis; 1971.

32. Hazelton P. and Murphy B. 2007. Interpreting soil test results: What do all the numbers mean? 2^nd^ Edition. CSIRO Publishing. 152p. 2007.

33. United States Department of Agriculture. Soil Mechanics Level 1 Module 3 USDA Soil Textural Classification Study Guide. USDA Soil Conservation Service, Washington DC; 1987.

34. Tekalign T. Soil, plant, water, fertilizer, animal manure and compost analysis. Working Document No. 13. International Livestock Research Center for Africa (ILCA), Addis Ababa, Ethiopia; 1991.

35. Olsen S., Cole C., Watanabe F. and Dean L. Estimation of available phosphorus in soils by extraction with sodium carbonate. USDA Circular 1954; 939, 1–19.

36. Landon J. Booker tropical soil manual: A handbook for soil survey and agricultural land evaluation in the tropics and subtropics. Longman Sci. and Tech. Publ., Harlow, UK; 1991.

37. FNRRC (Fogera National Rice Research and Training Center). Recommendation of Fertilizer Application for Practical Field Report, Woreta, Ethiopia; 2018.

38. Legesse D., Gezahegn A. and Tesfaye Z. Agricultural Technology Generation: Implications for Poverty Reduction. Paper presented on Ethel-Forum, Conference on Poverty Reduction, January 16-19 2002. Organized by Ethiopian Social Rehabilitation Fund. Addis Ababa. Ethiopia; 2002.

39. Derebe B., Bitew Y., Asargew F, Chakelie G, Optimizing time and split application of nitrogen fertilizer to harness grain yield and quality of bread wheat (Triticum Aestivum L.) in northwestern Ethiopia. PLoS ONE 2022; 17(12):2–17.

40. Biru A. Agricultural Experiment Management Manual, Part III. Institute of Agricultural Research. Addis Abeba; 1979.

41. Ghanbari-Bonjar A. Intercropped wheat and bean as a low-input forage, PhD thesis. Wye College. Univ. London); 2000.

42. Willey R. W and Rao M. R. A competitive ratio for quantifying competition between intercrops. Experimental Agriculture1980; 16 (2): 117–125.

43. Zhang G. Zaibin Y. & Shuting D. Interspecific competitiveness affects the total biomass yield in an alfalfa and corn intercropping system. Field Crops Research 2014; 124, 66–73.

44. Mead R., Willey R.W. The concept of land equivalent ratio and advantages in yields from intercropping. Experimental Agriculture 1980; 16: 217–228.

45. Hiebsch C.K. and Macollam R.E. 1980. Area time equivalency ratio. A method for evaluating the productivity of intercrops. Agronomy Journal 1980; 75: 15–22.

46. De Wit C.T., Ennik G. C., van den Bergh J. P. and Sonneveld A. Competition and nonpersistency as factors affecting the composition of mixed crops and awards. In Proceedings 8th international grassland congress 1960 (pp. 6–6).

47. Singh K. K., Ali M. and Venkatesh M. S. Pulses in cropping systems. Technical Bulletin, IIPR, Kanpur, 47; 2009.

48. Jolliffe P. A. and Wanjau F.M. Competition and productivity in crop mixtures: Some properties of productive intercrops. The Journal of Agricultural Science 1999; 132 (4): 425–35.

49. Willey R. W. Intercropping-its importance and research needs. Part 2. Agronomy and research approaches (No. REP-2605. CIMMYT.); 1979

50. CIMMYT (International Maize and Wheat Improvement Center). From Agronomic Data to Farmer Recommendations: An Economic Training Manual. Revised Edition. Mexico, D.F. 1988,

51. SAS (Statistical Analysis Software, Institute Inc.). JMP^®^ 13 Basic Analyses. Cary, NC: SAS Institute Inc; 2016.

52. Isaac A., Oyebisi A.K., Kayode O.S, Adeniji S. and Mojisola A.S. Effects of spatial arrangement and population density on the growth and yield of sesame (sesamum indicum l.) in a sesame/maize intercrop. Journal of Agricultural Sciences 2020; 65 (4): 337–350

53. Banik P. and Bagchi D. K. Evaluation of rice (Oryza sativa) and legume intercropping in upland situation of Bihar plateau. The Indian Journal of Agricultural Sciences 2006; 64(6): 3–4

54. Yayeh B., Fetien A. Tadess D. Competition Indices of Intercropped Lupine (Lal) and Small Cereals in Additive Series in West Gojam, North Western Ethiopia. American Journal of Plant Science 2014;, 5: 1296–1305

55. Chen C., Westcott K. M, Neill D. Wichman & Knox M. Row configuration and nitrogen application for barley-pea intercropping in Motana. Agronomy Journal 2004; 96: 1730–1738.

56. Egbe O. M., Alibo S. E. and Nwueze I. Evaluation of some extra-early-and early-maturing cowpea varieties for intercropping with maize in southern Guinea Savanna of Nigeria. Agriculture and Biology Journal of North America 2010 1(5): 845–858.

57. Jensen E. S., Peoples M. B., Boddey R. M., Brushoff P. M., Hauggaard-Nielsen H., J. R and Alves M. J. Legumes for mitigation of climate change and the provision of feedstock for biofuels and bio refineries. A review. Agronomy for sustainable development 2012; 32(2): 329–364.

58. Haymes R. and Lee H. C. Competition between autumn and spring planted grain intercrops of wheat (Triticum aestivum. L) And field bean (Vicia faba). Field crops research 1999; 62(2-3): 167–176.

59. Li L., Yang S., Li. X., Zhang F. and Christie P. Interspecific complementary and competitive interactions between intercropped maize and faba bean. Plant and soil 1999; 212(2): 105–114.

60. Vandermeer J. The ecology of intercropping, Cambridge Univ. Press. Cambridge. UK; 1989.

61. Reddy S. R., Reddy S. R., Reddi C. H., Reddy T. Y. and Reddi C. H. Principles of agronomy (Vol. 10). Kalyani publishers; 2007

62. Ghosh P. K. 2004. Growth, yield, competition and economics of groundnut/cereal fodder intercropping systems in the semi-arid tropics of India. Field crops research 2007’; 88(2-3): 227–237.

63. Trydemanknudsen M. H., Hauggaard-Nielsen H. B., Jornsgard B. and Jensen E.S. Comparison of inter-specific competition and N use in pea–barley, faba bean–barley, and lupine–barley intercrops grown at two temperate locations. The Journal of Agricultural Science2004; 142, 617–27.

64. Hauggaard-Nielsen H., Ambus P. and Jensen E. S. Interspecific competition, N use and interference with weeds in pea–barley intercropping. Field crops research 2001; 70(2), 101–109.

65. Tenaw W., Legesse H. and Gobeze L. Grain yield and economic benefit of intercropping barley and faba bean in the Highlands of Southern Ethiopia. East African Journal of Sciences 2016; 10(2), 103–110.

